# Astrocytes Regulate the Development and Maturation of Retinal Ganglion Cells Derived from Human Pluripotent Stem Cells

**DOI:** 10.1101/428631

**Authors:** Kirstin B. Langer, Ridhima Vij, Sarah K. Ohlemacher, Akshayalakshmi Sridhar, Clarisse M. Fligor, Elyse M. Feder, Michael C. Edler, Anthony J. Baucum, Theodore R. Cummins, Jason S. Meyer

## Abstract

Retinal ganglion cells (RGCs) form the connection between the eye and the brain, with this connectivity disrupted in numerous blinding disorders. Previous studies have demonstrated the ability to derive RGCs from hPSCs; however these cells exhibited some characteristics that indicated a limited state of maturation. Among the many factors known to influence RGC development in the retina, astrocytes are known to play a significant role in their functional maturation. Thus, efforts of the current study examined the functional maturation of hPSC-derived RGCs, including the ability of astrocytes to modulate this developmental timeline. Morphological and functional properties of RGCs were found to increase over time, with astrocytes significantly accelerating the functional maturation of hPSC-derived RGCs. The results of this study are the first of its kind to extensively study the functional and morphological maturation of RGCs in vitro, including the effects of astrocytes upon the maturation of hPSC-derived RGCs.

## Introduction

Human pluripotent stem cells (hPSCs) have emerged as an advantageous in vitro model for studying neural development and neurodegeneration, as these cells can be differentiated into any cell type of the body, including those of the nervous system (Corti et al., 2015; Jin et al., 2011; Marchetto et al., 2011; Meyer et al., 2009; Takahashi et al., 2007; Yu et al., 2007). More so, when derived from patient-specific sources, hPSCs can serve as powerful tools for disease modeling as well as large-scale pharmacological screens (Ebert and Svendsen, 2010; Grskovic et al., 2011; Inoue and Yamanaka, 2011). However, in order for hPSCs to effectively serve in these applications, they must closely mirror the phenotypic and functional features of the affected cell type. Retinal ganglion cells (RGCs) form the essential connection between the eye and the brain, allowing for visual transduction and the ability to see. This important connection becomes severed in numerous blinding disorders termed optic neuropathies, which cause progressive degeneration and eventual death of RGCs in the retina (Almasieh et al., 2012; You et al., 2013).

Previous studies have demonstrated the ability to successfully differentiate RGCs from hPSCs in an efficient and reproducible manner (Fligor et al., 2018; Gill et al., 2016; Langer et al., 2018; Maekawa et al., 2016; Ohlemacher et al., 2016; Riazifar et al., 2014; Sluch et al., 2015; Tanaka et al., 2015; Teotia et al., 2017). RGCs derived using such methods have recapitulated the developmental timing of the retina, as well as displayed various molecular and morphological characteristics similar to the in vivo cell type. As the projection neuron of the retina, RGCs extend long axons through the optic nerve in order to communicate visual information to the brain. hPSC-derived RGCs have demonstrated some capacity to exhibit predicted electrophysiological characteristics including ability to fire action potentials (APs), conduct ionic currents, and respond to glutamate (Gill et al., 2016; Ohlemacher et al., 2016; Riazifar et al., 2014; Sluch et al., 2015; Tanaka et al., 2015; Teotia et al., 2017). However, these studies have often demonstrated somewhat limited functional properties of hPSC-derived RGCs and have failed to take into consideration the milieu of factors known to influence RGC development and maturation. Among the more influential of these factors are contributions from astrocytes, which are known to closely associate with RGCs in the nerve fiber layer and optic nerve (Bussow, 1980; Vecino et al., 2016; Zuchero and Barres, 2015).

As such, the current study focused on characterizing the functional maturation of hPSC-derived RGCs based upon morphological complexity and electrophysiological properties, both over time as well as in response to signaling from astrocytes. Initially, hPSC-derived RGCs demonstrated increased neurite complexity as a function of time, including enlargement of the cell somas and elongation of neurites. Additionally, these hPSC-derived RGCs displayed increasing ionic currents, AP excitability, and the expression of synaptic proteins. To study the influence of astrocytes upon hPSC-derived RGC maturation, cells were either plated in co-culture with hPSC-derived astrocytes or supplemented with astrocyte conditioned media (ACM) to determine substrate-bound or secreted effects of astrocytes on RGCs, respectively. RGCs co-cultured with astrocytes demonstrated expedited and enhanced morphological and functional characteristics compared to RGCs cultured alone. The results of this study are the first of its kind to extensively study the functional and morphological maturation of RGCs in vitro as well as explore how astrocytes modulate and expedite this maturation. These results will also lead to the development of more realistic hPSC-derived disease models, with important implications for pharmacological screenings and cell replacement strategies in future studies.

## Results

### Increased RGC neurite complexity over time

RGCs exhibit complex morphological features within the retina, with long axons extending out of the eye through the optic nerve, while a complex and extensive network of dendrites reaches into the inner plexiform layer (Erskine and Herrera, 2014; Masland, 2012; Sernagor et al., 2001). Given this morphological complexity, an hPSC-derived RGC population should also exhibit intricate features in order to serve as an effective in vitro model. However, the study of hPSC-derived RGC morphological features has been somewhat limited in previous studies, as RGCs are often differentiated and examined in a mixed population of other cell types (Lamba et al., 2010; Meyer et al., 2011; Sridhar et al., 2013). Additionally, the definitive identification of RGCs within such cultures has been complicated by the lack of unique, reliable markers that allow for the visualization of entire RGCs including their extensive neurites. Thus, to overcome these issues, a CRISPR/Cas9-engineered cell line with a tdTomato reporter and mouse Thy1.2 selectable marker was used to identify and enrich for hPSC-derived RGCs (Sluch et al., 2017).

In accordance with previous studies, hPSC-derived RGCs could be readily observed within the first 40 days of differentiation (Ohlemacher et al., 2016), at which point RGCs were enriched and plated to allow for maturation and examination. Over the next 10 days, hPSC-derived RGCs displayed features of increasing neurite complexity, including robust and complex neurite outgrowth (Figure 1). Purified RGCs demonstrated a dramatic and significant increase in soma size, more than doubling within the first 10 days of growth (Figure 1J). Additionally, RGC neurites grew significantly in length over time, with neurites extending upwards of 150μm within 10 days of plating (Figure 1F-I, K). The complexity of RGC neurites was examined and displayed increasingly complex morphologies, with the neurite branch order more than doubling within this time course (Figure 1L). RGC neurites also displayed a 13-fold increase in the number of branch points within the first 10 days of growth (Figure 1M). As such, hPSC-derived RGCs were capable of exhibiting increased morphological complexity over time, including the enlargement of the cell soma and elongation of complex neurites.

**Figure 1:**
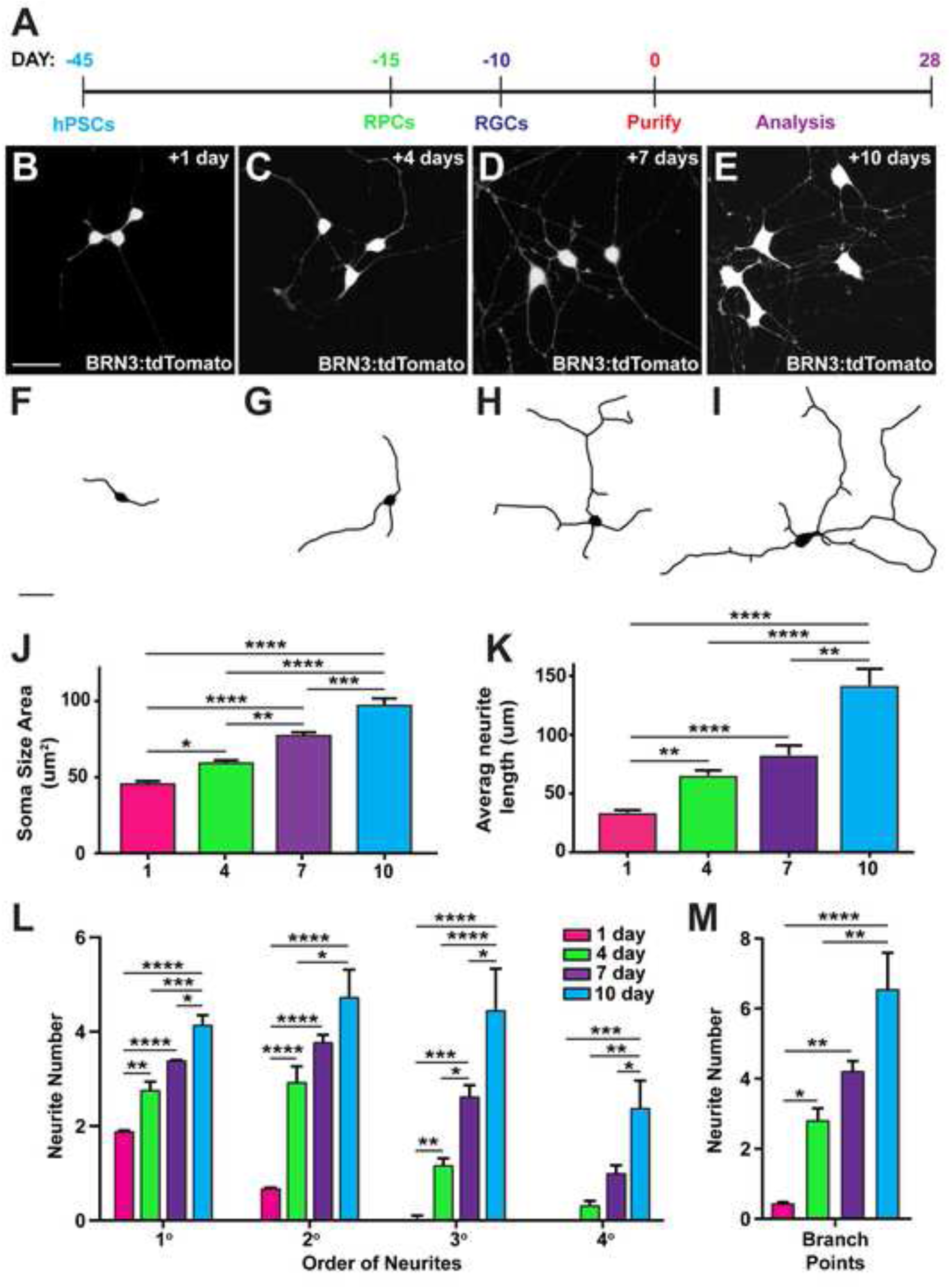
Temporal acquisition of complex neuronal morphologies by hPSC-derived RGCs. (A) Representative timeline for the experimental differentiation and analysis of hPSC-derived RGCs. (B-I) Immunocytochemistry and representative tracings of hPSC-derived RGCs at 1- (B,F), 4- (C,G), 7- (D,H), and 10-days (E,I) of differentiation displayed enhanced morphological maturation over time. (J) The area of RGC somas significantly increased over time. (K) The average neurite length of hPSC-derived RGCs significantly increased from 1 to 10 days of differentiation. (L,M) Lastly, neurite tracings demonstrated a significant increase in neurite order complexity (L) and number of branches (M) within 10 days of differentiation. Scale bars equal 30μm. Error bars represent SEM (n=5) and significances were determined using a 95% confidence interval; *<0.05, **<0.01, ***<0.001, ****<0.0001.

### Functional maturation of hPSC-derived RGCs over time

Within the retina, RGCs are the most excitable cells, with the nearly exclusive ability to fire APs necessary for the propagation of visual signals to the brain (Erskine and Herrera, 2014; Gao and Wu, 1999; Guenther et al., 1999; Meister et al., 1991; Sernagor et al., 2001; Tian et al., 1998; Wang et al., 1997). As such, efforts were undertaken to examine the electrophysiological features of hPSC-derived RGCs, as well as how these characteristics change over time (Figure 2). Voltage-clamp recordings from hPSC-derived RGCs exhibited an increase in peak sodium conductance as neurons matured from 1 to 4 weeks (Figure 2A). The current-voltage relationship (I-V curve; Figure 2B) displayed increased sodium currents with the average peak current increasing over time. The voltage dependence of activation for normalized peak sodium conductance showed a significant hyperpolarized shift as the RGCs matured from week 1 to week 2 and further (Figure 2C), signifying functional maturation of hPSC-derived RGCs that is consistent with the developmental shift observed in rat and mice RGCs (Rothe et al., 1999; Schmid and Guenther, 1998). Current density plots also displayed an increase in sodium currents from 1 to 4 weeks of RGC maturation, indicating an increase in the expression of voltage-gated sodium channels (Figure 2D).

**Figure 2:**
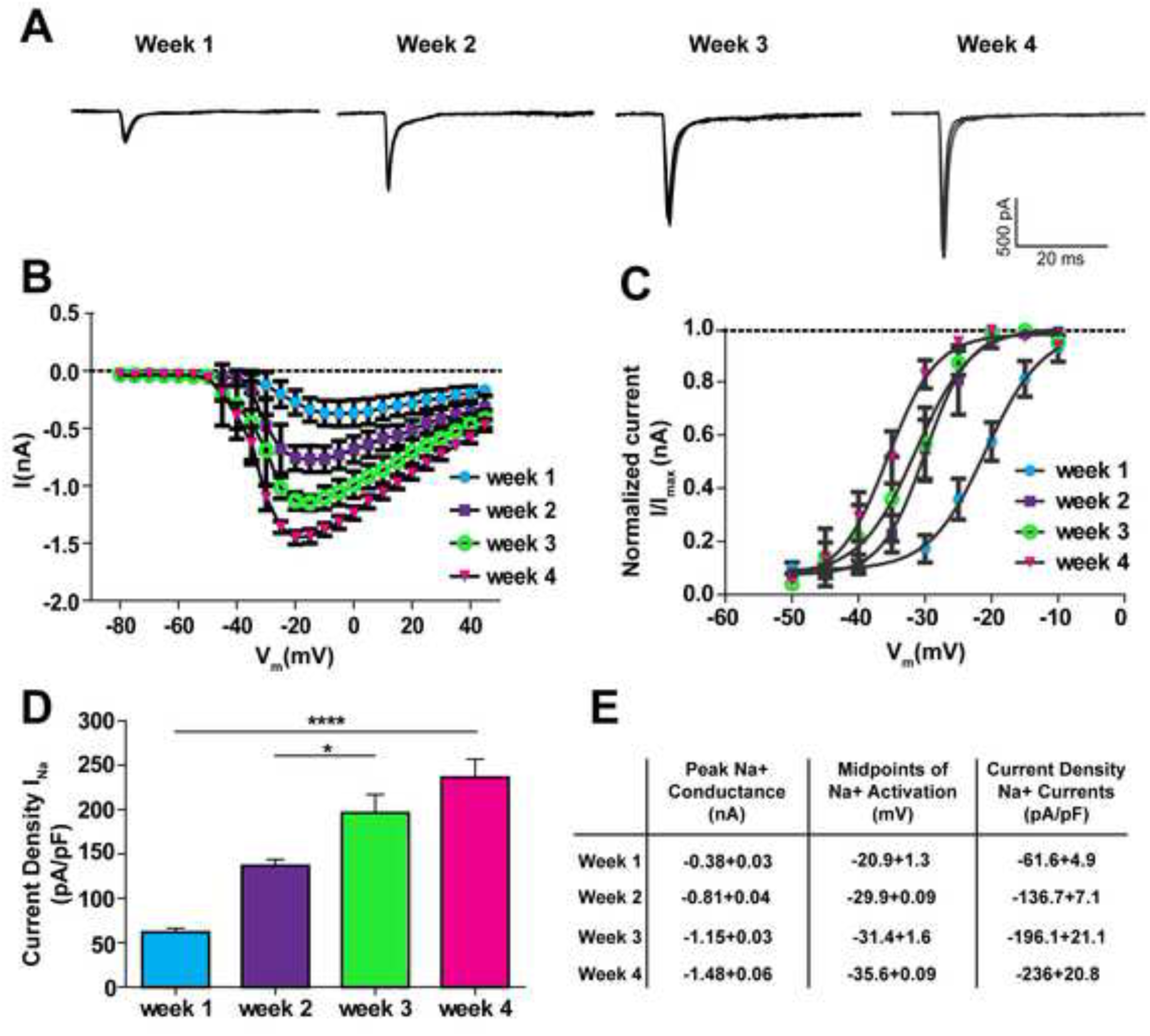
Temporal development of functional maturation of hPSC-derived RGCs. (A) Patch-clamp recordings revealed an increase of inward ionic currents within 4 weeks of differentiation. (B) From 1 to 4 weeks, IV-curves displayed increased inward current as RGCs matured. (C) Voltage dependence of activation indicated a significant shift in the hyperpolarized direction over time. (D) A current density plot demonstrated a significant increase in sodium currents over 4 weeks. (E) A table representing the increases in peak conductance, midpoints of activation, and current density values at each time point. Error bars represent SEM (n > 4 for each timepoint) and significant differences indicated as *<0.05, ****<0.0001.

Subsequently, the ability of hPSC-derived RGCs to fire spontaneous APs was examined, with no spontaneous activity detected after 1 week, but with spontaneous APs elicited as early as 2 weeks of differentiation (Figure 3A). By 4 weeks, repetitive AP firing patterns emerged, along with occasional AP bursts. The percentage of hPSC-derived RGCs capable of spontaneous AP firing was found to continually increase from 1 to 4 weeks of differentiation (Figure 3B). Furthermore, the average number of spontaneous APs fired increased significantly as hPSC-derived RGCs matured (Figure 3C).

**Figure 3:**
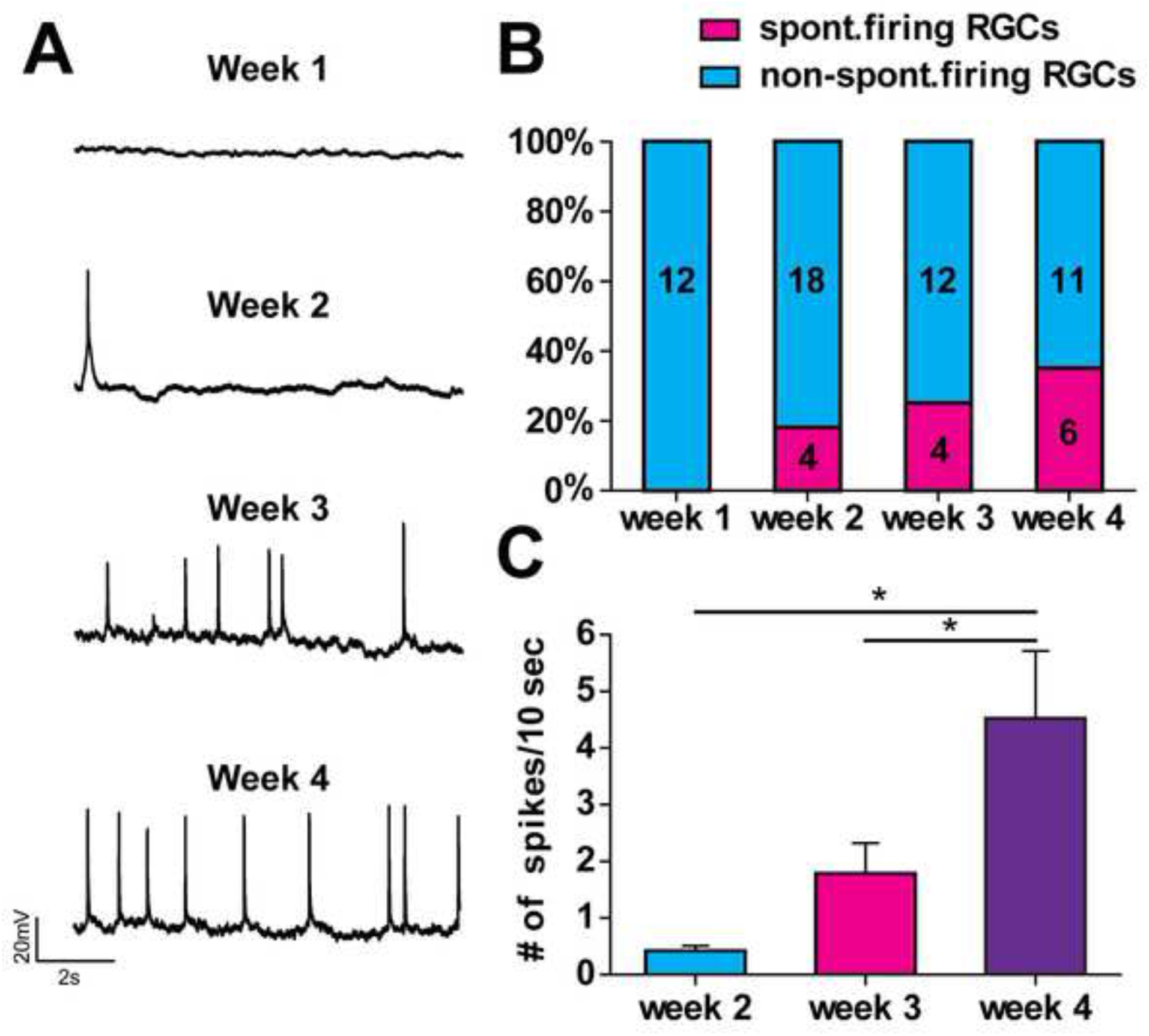
Increased excitability of hPSC-derived RGCs in a temporally-dependent manner. (A) Patch-clamp recordings demonstrated the increased ability of hPSC-derived RGCs to fire spontaneous APs from 1 to 4 weeks of differentiation. (B) The frequency of spontaneous APs increased between 1 (n=12), 2 (n=22), 3 (n=16), and 4 (n=17) weeks. (C) Spiking behavior of hPSC-derived RGCs from 1 to 4 weeks significantly increased. Error bars represent SEM and significant differences indicated as *<0.05.

To determine if hPSC-derived RGCs were capable of forming presumptive synaptic contacts, the expression of various synaptic proteins was analyzed from 1 to 3 weeks of differentiation (Figure 4). hPSC-derived RGCs demonstrated a significant increase in the number of SV2-positive puncta along their neurites from 1 to 3 weeks (Figure 4 A-D). Additionally, western blot analyses revealed an increased expression of synaptic proteins over the course of differentiation, including Synapsin-1, vGlut2, Neurexin-1, and SNAP25 (Figure 4E-F). Taken together, these data reveal that hPSC-derived RGCs exhibited electrophysiological properties and formed presumptive synaptic contacts as predicted by in vivo studies, with these features increasing over the course of differentiation.

**Figure 4:**
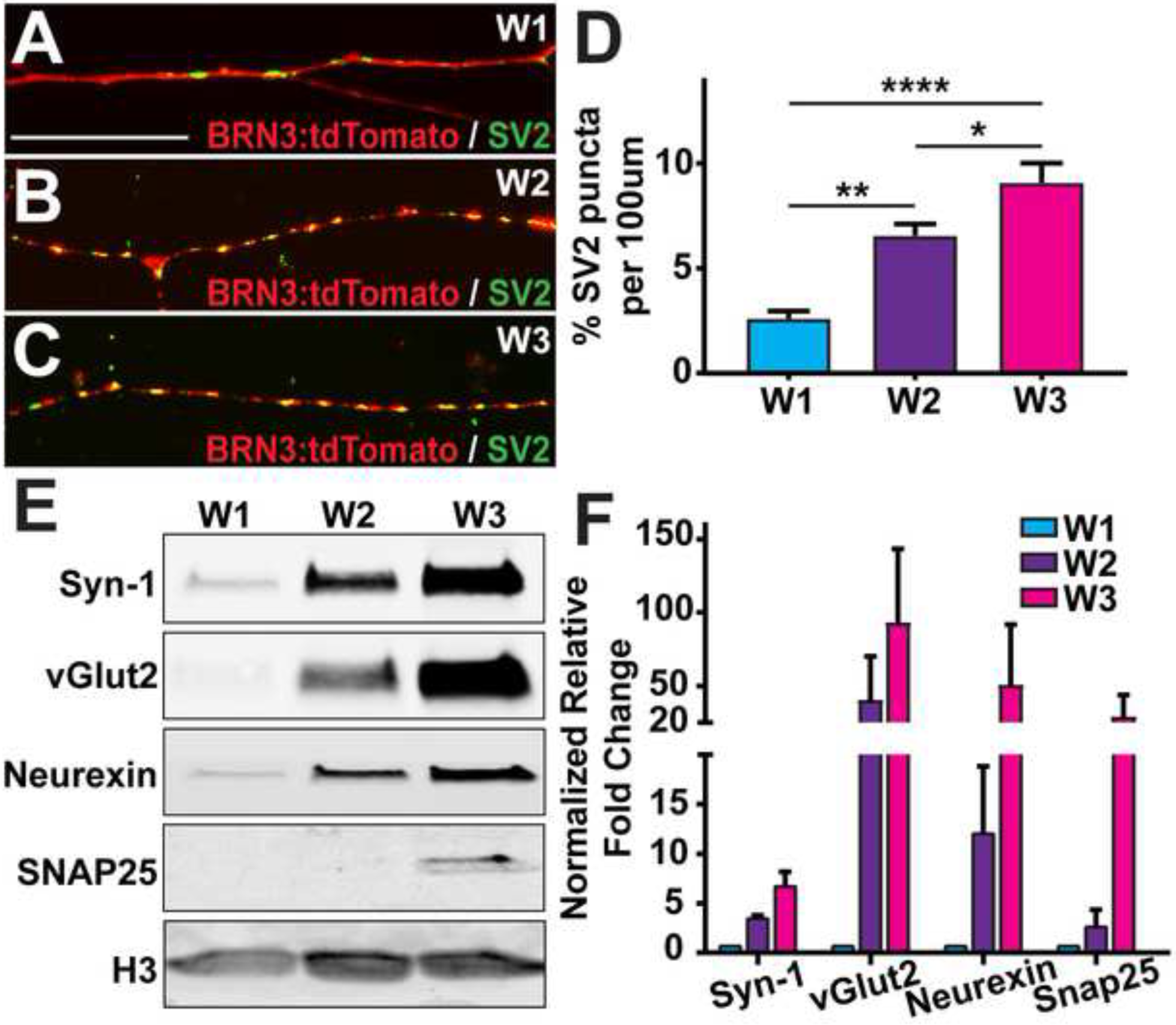
Formation of presumptive synaptic contacts by hPSC-derived RGCs. (A-C) Immunocytochemistry demonstrated the increased presence of SV2-puncta within hPSC-derived RGC neurites at 1 (A), 2 (B), and 3 (C) weeks of differentiation. (D) The number of SV2-puncta per neurite length significantly increased over time. (E-F) Western blot analyses displayed an increased expression levels of a number of synaptic proteins. Scale bars equal 20μm. Error bars represent SEM (n=3) and significant differences were determined at a 95% confidence interval; *<0.05, **<0.01, ***<0.001, ****<0.0001.

### Astrocytes modulate the morphological complexity of hPSC-derived RGCs

Within the retina, astrocytes are closely associated with RGCs within the nerve fiber layer and optic nerve, where they are known to support RGCs through the release of growth factors, reuptake of neurotransmitter, and maintaining ionic balances, among other roles (Allen and Barres, 2009; Barres, 1991; Ogden, 1978; Pellerin and Magistretti, 1994; Vecino et al., 2016; Wordinger et al., 2003). Additionally, astrocytes have been shown to modulate the maturation of a variety of neuronal cell types and influence their synaptic development in vitro (Johnson et al., 2007; Odawara et al., 2014). However, similar experiments have not tested their ability to influence RGC development and maturation. As such, co-cultures with hPSC-derived astrocytes were developed as a strategy to enhance the maturation of hPSC-derived RGCs.

hPSCs were differentiated into an astrocytic lineage (Krencik and Zhang, 2011), yielding highly enriched populations of GFAP-positive astrocytes after 9 months of growth (Supplemental Figure 1). Additionally, these astrocyte populations also expressed a multitude of other glial cell markers, including Vimentin, S100β, SOX2, and Nestin (Supplemental Figure 1C-E). The ability of these hPSC-derived astrocytes to support RGCs was also suggested by the expression of Excitatory Amino Acid Transporter 1 (EAAT1), important for the reuptake of excess glutamate at the synapse (Supplemental Figure 1F). To establish a co-culture system with which to directly study the interactions between astrocytes and RGCs, hPSC-derived RGCs were purified and plated in association with hPSC-derived astrocytes and grown in co-culture for 10 days to observe changes in RGC morphology and neurite outgrowth (Supplemental Figure 1G). Additionally, to separate the effects of astrocytes as either substrate-bound or due to soluble factors secreted into the media, hPSC-derived RGCs were separately supplemented with astrocyte conditioned media (ACM).

After 10 days of plating, purified RGCs co-cultured with astrocytes displayed significant and robust neurite outgrowth and complexity compared to that of RGCs grown alone or supplemented with ACM (Figure 5A-F). Astrocyte co-cultures led to significant increases in RGC soma size as well as the average length of the longest neurite, while ACM showed no significant effects (Figure 5G-H). More so, RGCs co-cultured with astrocytes displayed significantly more complex neurite outgrowth as indicated by the number of primary, secondary, tertiary, and quaternary neurites and number of branch points (Figure 5I-J). As supplementing RGCs with ACM led to no significant changes in morphological analyses, subsequent experiments examined the effects of astrocytes solely through direct contact in co-cultures.

**Figure 5:**
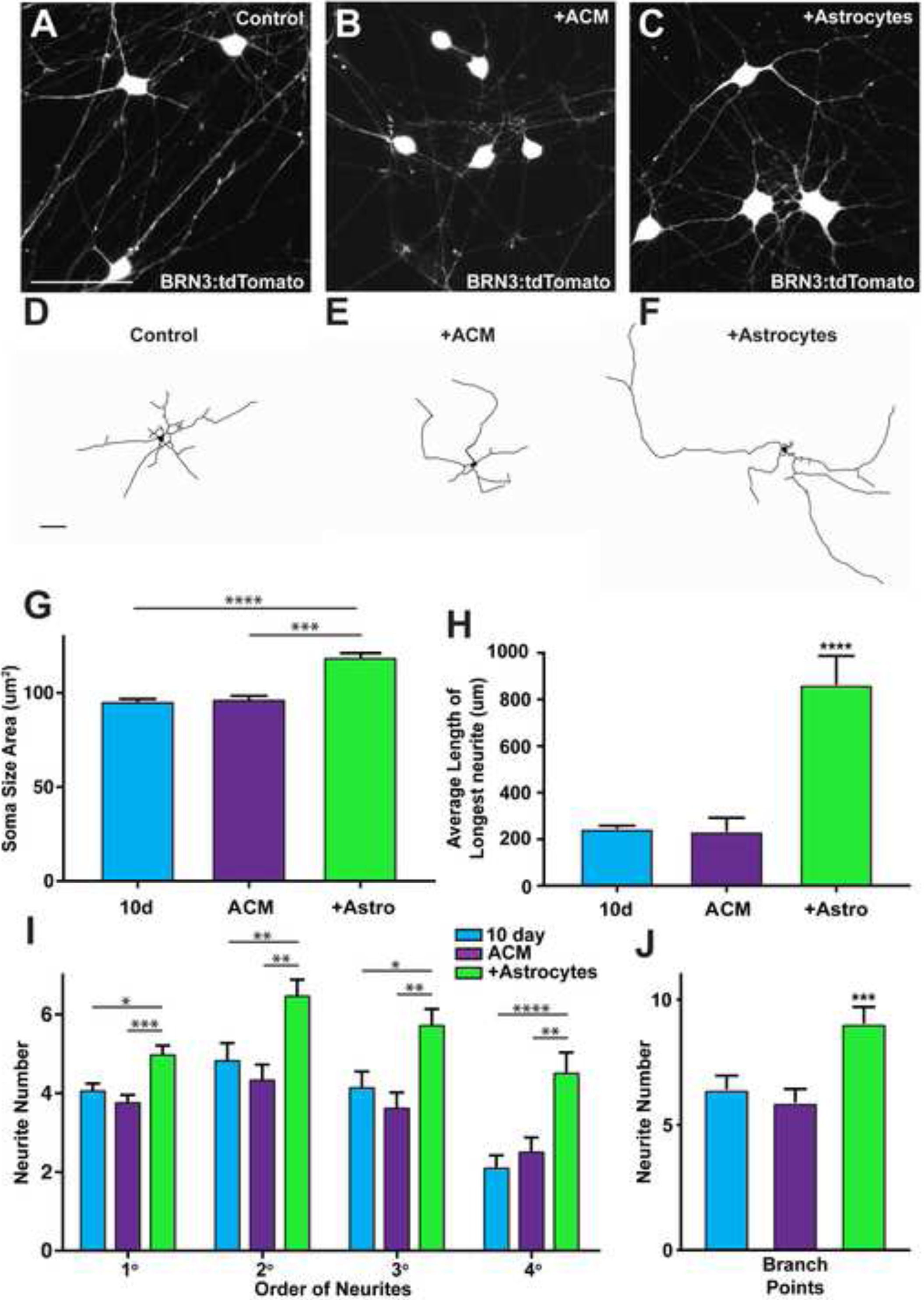
Astrocytes expedite and enhance the morphological maturation of hPSC-derived RGCS. (A-F) At 10 days of differentiation, immunocytochemistry and representative neurite tracings of hPSC-derived RGCs displayed enhanced morphological maturation for RGCs grown with astrocytes (C, F) compared to RGCs grown alone (A,D) or RGCs supplemented with ACM (B, E). (G-H) Significant increases were observed in soma size area (G) and the length of the longest neurite (H) for hPSC-derived RGCs grown with astrocytes. (I-J) Neurite tracings displayed a significant increase in neurite order complexity (I) and number of branches (J) for hPSC-derived RGCs on astrocytes compared to other conditions. Scale bars equal 50μm. Error bars represent SEM (n=5) and significant differences were determined at a 95% confidence interval; *<0.05, **<0.01, ***<0.001, ****<0.0001.

### Astrocytes accelerate the functional maturation of hPSC-derived RGCs

Next, the effects of astrocyte co-cultures on the functional maturation of hPSC-derived RGCs were characterized. Electrophysiological analyses of the evoked APs were conducted both on the RGCs grown alone and those co-cultured with astrocytes at 3 weeks of differentiation (Figure 6A-D). Quantitatively, RGCs co-cultured with astrocytes demonstrated an increased number of AP firings (Figure 6E), both in response to a steady current injection as well as in response to a ramp depolarization event. Additionally, the presence of astrocytes in co-culture with hPSC-derived RGCs increased the AP amplitude (Figure 6F), and decreased the AP duration (Figure 6G) when compared to RGCs cultured alone. Furthermore, RGCs co-cultured with astrocytes showed a more hyperpolarized AP threshold (Fig. 6H).

**Figure 6:**
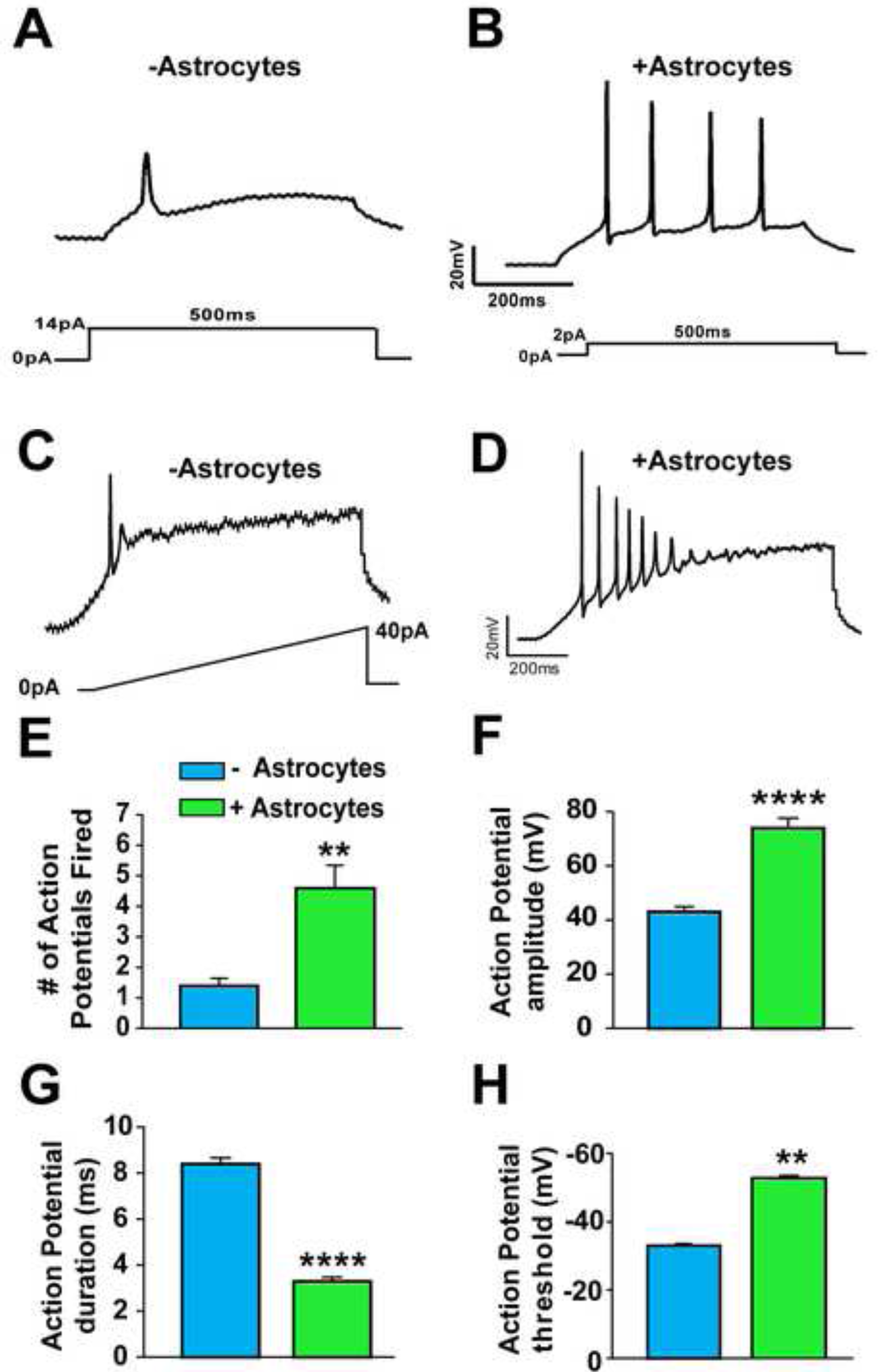
hPSC-derived astrocytes modulate the functional maturation of hPSC-derived RGCs. (A-B) Patch-clamp recordings demonstrated the heightened excitability of hPSC-derived RGCs grown on astrocytes (B) compared to hPSC-derived RGCs grown alone (A). (C-D) A ramp depolarization indicated the ability for hPSC-derived RGCs grown with astrocytes (D) to fire more and repetitive APs compared to RGCs alone (C). (E-H) hPSC-derived RGCs grown with astrocytes exhibited a higher number of APs fired (E), increased amplitude (F), decreased duration (G) and a more hyperpolarized AP threshold (H). Error bars represent SEM and significances were determined at a 95% confidence interval; **<0.01, ****<0.0001.

Additionally, the ability of astrocytes to enhance the formation of presumptive synaptic contacts was assessed by comparing RGCs grown alone to RGCs co-cultured with astrocytes. RGCs grown with astrocytes displayed a significantly higher number of SV2-positive presumptive synaptic puncta (Figure 7A-C). Analyses of spontaneous synaptic currents revealed that RGCs co-cultured with astrocytes also exhibited more frequent spontaneous inward ionic currents (Figure 7D). Furthermore, western blot analyses revealed that RGCs grown in co-culture with astrocytes demonstrated significantly increased expression of a variety of synaptic proteins (Figure 7E-F). Additional western blot analysis confirmed that increased synaptic protein expression in co-cultures was a result of increased RGC maturation and not derived from astrocyte populations (Supplemental Figure 2).

**Figure 7:**
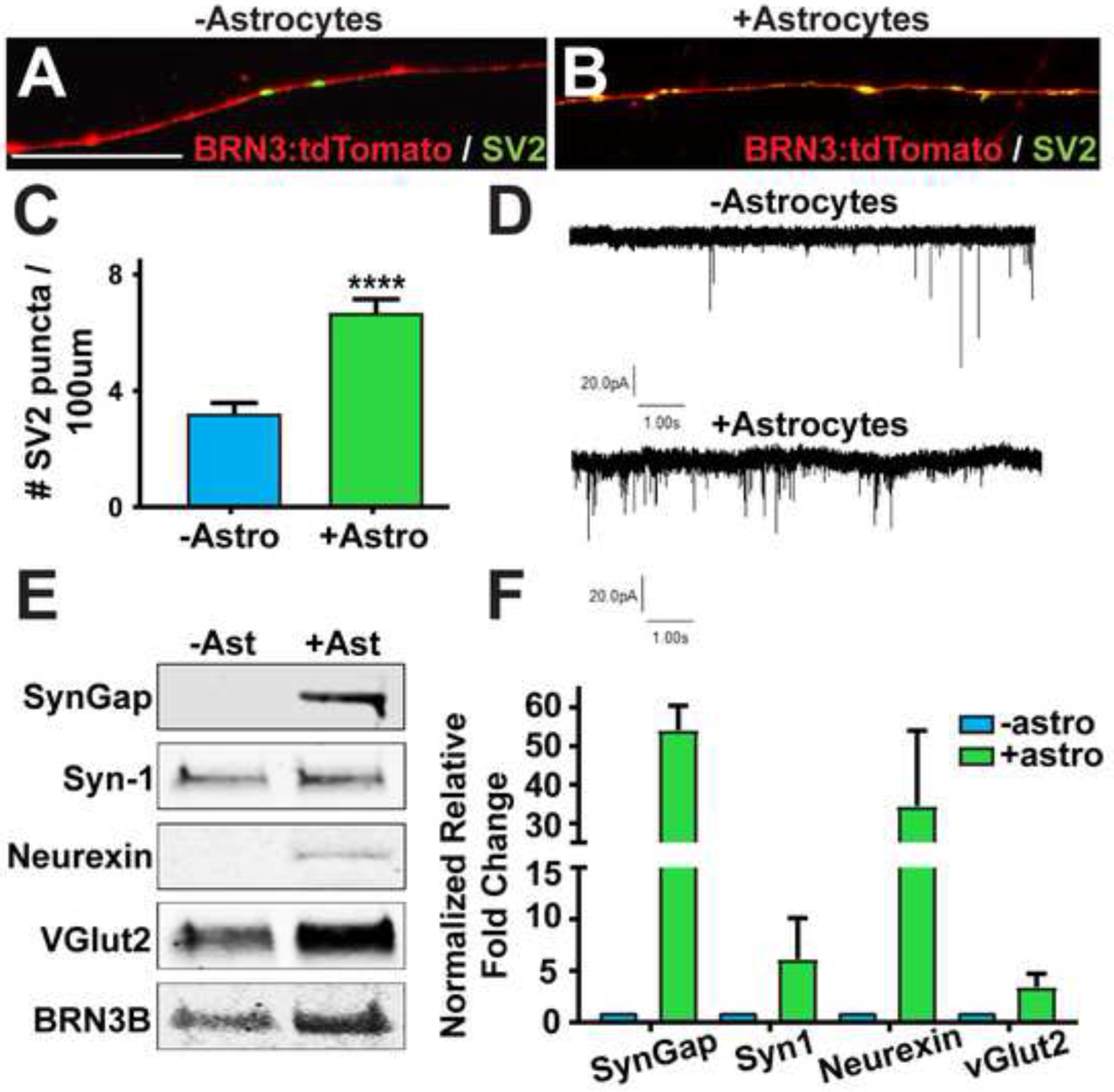
Astrocytes regulate the synaptic maturation of hPSC-derived RGCs. (A-B) At 10 days of differentiation, immunocytochemistry demonstrated the increased presence of SV2-puncta within hPSC-derived RGC neurites grown in the presence of astrocytes compared to those grown alone. (C) The number of SV2 puncta significantly increased in hPSC-derived RGCs grown with astrocytes. (D) Patch-clamp recordings demonstrated an increase in the number and frequency of spontaneous postsynaptic currents for RGCs grown with astrocytes. (E-F) Western blot analyses demonstrated an increase in the expression of a number of synaptic proteins when hPSC-derived RGCs were grown with astrocytes. Scale bars equal 20μm. Error bars represent SEM (n=3) and significances were determined at a 95% confidence interval; ****<0.0001.

## Discussion

The utilization of hPSCs to study RGCs has been more recently studied with an important emphasis on how these cells can serve as a translational model for both the development of RGCs as well as diseases resulting in degeneration and eventual death of these cells (Corti et al., 2015; Gaillard and Sauve, 2007; Jin et al., 2011). However, to date, hPSC-derived RGCs have often exhibited limited functional capabilities and have overlooked the interaction of many other cell types closely associated with RGCs in the retina that can play a vital role in the development and maturation of these cells (Bussow, 1980; Vecino et al., 2016). As RGCs form the essential connection between the eye and the brain, these cells demonstrate a number of morphological and functional characteristics unique to RGCs within the retina, including the extension of long and complex neurites and the ability to fire APs (Guenther et al., 1999; Hebel and Hollander, 1983; Sernagor et al., 2001; Wang et al., 1997). The generation of hPSC-derived RGCs which recapitulate these distinctive characteristics would allow for a more representative model of these cells, with important implications for how these cells can serve as a tool for disease modeling, drug screenings and future cell replacement therapies. The results of this study demonstrate the derivation of RGCs that exhibit characteristics closely resembling their in vivo cell counterpart, allowing for a greater understanding of how these cells develop and may be affected in disease states. Furthermore, the ability to extensively study the morphological and functional maturation of hPSC-derived RGCs over time demonstrates their ability to acquire advancing features of RGC maturation, with astrocytes significantly expediting and enhancing this maturation.

RGCs are the main cell type affected in numerous blinding disorders which result in the degeneration and death of these cells leading to loss of vision and eventual blindness (Almasieh et al., 2012; Quigley, 2011; You et al., 2013). Therefore, the use hPSC-derived RGCs allows for the establishment of translational models of these diseases to understand mechanisms causing the death of RGCs, as well as the development of treatments and therapeutics targeting these disease phenotypes, including large scale pharmaceutical screenings and potential cell replacements strategies. However, in order for hPSC-derived RGCs to serve as a reliable disease model, they must closely mimic the phenotypic and functional characteristics of the affected cell type, as disease states often manifest in fully differentiated and functionally mature cells. In the current study, hPSC-derived RGCs exhibited increased morphological complexity and functional maturation over time, including increasing inward ionic currents and the more frequent firing of spontaneous APs, with these features becoming significantly enhanced in the presence of astrocytes. The demonstration of these advancing features of RGC maturation thus allows for the ability to use these cells as a more faithful model for studying diseases affecting the heath of RGCs, as well as aid in the development of therapeutic strategies to target disease mechanisms.

A number of previous studies have demonstrated the ability to derive RGCs from hPSCs, with several of these studies establishing that these RGCs may possess a range of functional properties including ionic currents, firing of APs, and response to glutamate (Gill et al., 2016; Ohlemacher et al., 2016; Riazifar et al., 2014; Sluch et al., 2015; Tanaka et al., 2015; Teotia et al., 2017). However, these results often indicated a degree of immaturity of these cells, and seldom examined their functional properties beyond a single time point post-differentiation. Results of the current study observed hPSC-derived RGCs over multiple time points, demonstrating progressive features of their development and growth as they differentiated through various stages of morphological and functional maturity. By three weeks of differentiation, hPSC-derived RGCs were capable of conducting substantial inward ionic currents with a hyperpolarized voltage-dependence of activation, firing trains of spontaneous APs, and expressing a variety of synaptic proteins, all characteristics consistent with a functionally mature RGC. The study of hPSC-derived RGCs in such a manner allows for their use as a more appropriate developmental model.

RGCs are the projection neuron of the retina forming the vital connection between the eye and the brain, doing so by extending long and intricate axons through the optic nerve as well as forming complex connections with retinal interneurons within the inner plexiform layers (Masland, 2012). The analysis of various morphological properties within 10 days of differentiation demonstrated a significant increase in the length and complexity of hPSC-derived RGCs neurites over time. Similarly, hPSC-derived RGCs somas were shown to significantly increase in area over time, consistent with RGCs found within the retina that often possess the largest cell bodies. Taken together, these results indicated the ability for hPSC-derived RGCs to acquire advancing features of morphological maturation and as such, establish hPSC-derived RGCs as a more reliable model of RGC development.

RGCs form synaptic contacts both in the retina and the brain, with pre-synaptic contacts formed with bipolar and amacrine cells in the inner plexiform layer, and post-synaptic contacts formed with targets including the lateral geniculate nucleus and the superior colliculus (Herrera et al., 2017; Martersteck et al., 2017). The presence of synaptic features within hPSC-derived RGCs would thus serve as an additional measure of their maturation and likewise, their effectiveness as an in vitro model. A number of synaptic proteins were found to be expressed by hPSC-derived RGCs over time, indicating the formation of presumptive synaptic contacts. The expression of these proteins was found to increase over time, indicating the acquisition of advancing features of maturation. The prospective ability of these hPSC-derived RGCs not only contributes to their effectiveness as an appropriate in vitro model, but may also enable their use for cellular replacement if they are capable of incorporating into a neural network.

While hPSC-derived RGCs exhibited numerous features of maturation, many other cell types are known to play a role in this process in vivo. Astrocytes are found in close association with RGCs in the nerve fiber layer of the retina as well as within the optic nerve (Vecino et al., 2016). These cells provide support to RGCs in a number of ways, including the regulation ionic concentrations, the reuptake of excess neurotransmitters, and the release of various growth factors (Clarke and Barres, 2013; Eroglu and Barres, 2010; Pellerin and Magistretti, 1994; Pfrieger and Barres, 1997; Ullian et al., 2004). Therefore, the interaction of these two cell types could enhance the maturation of hPSC-derived RGCs when grown in a co-culture environment that was more similar to the retina. In the current study, the manner in which astrocytes could modulate RGC maturation was examined, with effects possibly due to the release of soluble factors or through direct contact between astrocytes and RGCs. hPSC-derived RGCs grown in direct contact with astrocytes displayed expedited and significantly enhanced morphological and functional maturation compared to RGCs grown alone or in the presence of astrocyte-derived soluble factors tested with ACM, including the extension of longer and more complex neurites as well as the ability to fire APs. As soluble factors tested by ACM did not demonstrate any significant differences compared to control experiments, the effects observed via direct contact with astrocytes could potentially be explained by the development of gap junctions between astrocytes and RGCs, or perhaps through the activation of signaling pathways triggered through substrate-bound ligands found on the astrocytic cell membrane. Regardless, these results established the importance of direct contact between hPSC-derived RGCs with hPSC-derived astrocytes, which more accurately represents an environment more similar to that found within the retina and optic nerve.

Astrocytes within the central nervous system are known to possess various regional identities depending on their location (Ben Haim and Rowitch, 2017; Hochstim et al., 2008; Tsai et al., 2012). For example, astrocytes within the retina are known to express the transcription factor PAX2 similar to those within the midbrain where retinal astrocytes originate (Chu et al., 2001; Scheef et al., 2005; Sehgal et al., 2009). Conversely, other transcription factors can identify the regionalization of astrocytes elsewhere in the central nervous system. Although the expression of markers within astrocytes may vary based upon their location, whether or not this heterogeneity results in functional variability remains unknown, with all astrocytes sharing the common function of supporting neurons through the regulation of homeostasis as well as aiding in synaptic formation and maturity (Clarke and Barres, 2013; Eroglu and Barres, 2010; Pellerin and Magistretti, 1994; Pfrieger and Barres, 1997; Ullian et al., 2004). While the astrocytes used within the current study did not express PAX2 similar to native retinal astrocytes, they did express a multitude of other astrocyte-associated markers and exhibited a significant effect on the maturation of hPSC-derived RGCs, as would be expected by retinal astrocytes.

One the most profound effects of astrocytes upon hPSC-derived RGCs was the enhancement of their electrophysiological properties. When grown with astrocytes, these cells had a higher level of activity, including increased firing of APs and presumptive synaptic currents. Additionally, these characteristics were often easier to elicit in the presence of astrocytes as indicated by the injection of smaller amounts of current, indicating a heightened excitability of hPSC-derived RGCs. Taken together, these results indicated that co-culturing astrocytes with hPSC-derived RGCs created an expedited and enhanced degree of maturation compared to RGCs cultured alone. More so, RGCs co-cultured with astrocytes generated a cell type more closely resembling in vivo RGCs with the ability to extend significantly longer and more intricate neurites, fire repetitive APs, and form presumptive synaptic contacts.

Overall, the results of the current study indicate the important contributions of astrocytes to RGC development and maturation. In future studies, the role of astrocytes should be considered in the establishment of developmental and disease models utilizing RGCs. The interaction of astrocytes with RGCs would allow for the development of a more mature cell type that can be used to properly identify and further understand disease deficits affecting RGCs. More so, given the important role of astrocytes in RGC maturation and function, future studies should also focus on the contributions of astrocytes to disease states and if diseased astrocytes are playing a role in the degeneration and death of RGCs. Furthermore, given their importance in the maturation and homeostasis of RGCs, astrocytes may provide an alternative target for therapeutics for optic neuropathies, and may also serve as a target for cell replacement strategies in future studies in order to enhance the survival and function of endogenous RGCs.

## Materials and Methods

### Maintenance of hPSCs

Human pluripotent stem cells (hPSC) were grown as previously described (Ohlemacher et al., 2015). Multiple hPSC-lines were used throughout this study including H7 (Thomson et al., 1998), H9 (Thomson et al., 1998), TiPS5 (Phillips et al., 2012), and miPS2 (Sridhar et al., 2016). In short, hPSCs were maintained in the undifferentiated state using hESC-qualified matrigel (Corning) on six-well plates with mTeSR1 medium (StemCell Technologies) at 37°C and 5% CO2. Passaging of hPSCs was accomplished by enzymatically lifting colonies using dispase (2mg/mL) for approximately 15 minutes. Colonies were then mechanically disrupted to yield small aggregates of cells and split at a 1:6 ratio.

### Differentiation of hPSCs into RGCs

hPSCs were differentiated to a retinal fate using previously described protocols with minor modifications (Ohlemacher et al., 2015). After 30 days of differentiation, early retinal organoids could be morphologically identified and isolated apart from other forebrain neurospheres. At day 45, retinal organoids were dissociated using Accutase for 20 minutes at 37°C. Following dissociation, single cells were immunopurified for RGCs with the Thy1.2 surface receptor using the MACS magnetic cell separation kit (Miltenyi Biotec). Briefly, dissociated cells were incubated with CD90.2 microbeads for 15 minutes at 4°C in the dark at a ratio of 10uL of beads: 90uL of MACs Rinsing Buffer per 10 million cells (Sluch et al., 2017). Purified RGCs were then plated on poly-d-ornithine and laminin-coated coverslips at a density of 10,000 cells/coverslip and maintained in BrainPhys Neuronal Media (StemCell Technologies) supplemented with 20ng/mL CNTF. Coverslips were then fixed at indicated timepoints post plating to analyze RGC maturation, or utilized for subsequent patch clamp recordings.

### Astrocyte Differentiation from hPSCs

hPSCs were differentiated to astrocytes using previously described protocols (Krencik and Zhang, 2011). Within 40 days of differentiation, early neurospheres were supplemented with 20 ng/ml EFH (FGF2 and EGF) to induce proliferation to a gliogenic phase. Neurospheres were chopped and expanded using a mechanical tissue chopper to a size of 200μm every 2 weeks for up to 12 months. Between 9 and 12 months, neurospheres became enriched for astrocytes and were then dissociated using Accutase for 20 minutes at 37°C. Single cells were plated at a density of 20,000 astrocytes per 12 mm coverslip and supplemented with BrainPhys Neuronal Media with the addition of 20ng/mL of CNTF to yield GFAP-positive astrocytes within 3 weeks of plating. Subsequently, co-cultures of RGCs and astrocytes were established with 10,000 purified RGCs plated with adhered astrocytes to create a substrate-bound co-culture. Additionally, astrocyte conditioned media (ACM) was collected from plated astrocytes, filtered to remove cellular components, and added every other day to RGC cultures at a dilution of 1:1 together with fresh BrainPhys as previously described (Pfrieger and Barres, 1997). Co-culture experiments were fixed at indicated time points post plating and analyzed for RGC maturation, or utilized for patch clamp analysis.

### Immunocytochemistry

Samples were fixed with 4% paraformaldehyde for 30 minutes and then washed 3 times with phosphate buffered saline (PBS). Cells were then permeabilized in 0.2% Triton X-100 for 10 minutes at room temperature, followed by blocking in 10% donkey serum for one hour at room temperature. Primary antibodies (Supplemental Table 1) were diluted in 5% donkey serum and 0.1% Triton X-100 solution and applied to samples overnight at 4°C. The following day, cells were washed 3 times with PBS and blocked with 10% donkey serum for 10 minutes at room temperature. Secondary antibodies were diluted in 5% donkey serum and 0.1% Triton X-100 solution and added for 1 hour at room temperature. Cells were then washed 3 times with PBS and mounted onto slides for microscopy. Immunofluorescent images were obtained using a Leica DM5500 fluorescence microscope.

### Quantification and Statistical Analyses

Samples were analyzed for RGC maturation based on soma size, neurite length and complexity. Numerous biological replicates (n=5) were obtained at each time point. Immunofluorescent images were taken and analyzed for the expression of tdTomato to identify RGCs. The RGC soma size area, neurite length, and branch order were quantified and recorded using Image J. Statistical analyses were performed using One-Way ANOVA followed by Tukey’s post hoc. Statistical differences were determined based on a *p* value of less than 0.05. Additionally, the abundance of SV2 puncta was quantified using Image J, with the total neurite length associated with SV2 immunoreactivity quantified using Image J plugin NeuronJ. Statistical analyses were performed using One-Way ANOVA followed by Tukey’s post hoc for SV2 expression in RGCs alone over time and a Student’s t-test for comparing SV2 expression in RGCs to those RGCs grown on astrocytes. Statistical differences were determined based on a *p* value of less than 0.05.

### Western Blot

RGCs were collected and sonicated in a 2% SDS solution. Protein concentration was determined using a BCA assay (ThermoScientific) and samples were normalized so that equal amounts of protein were loaded into each well. Lysates were mixed with 4x Sample Buffer plus DTT and loaded onto 4–15% gradient gels. Gels were transferred to nitrocellulose using the Trans-Blot Turbo system (BioRad), blocked with 5% milk in tris buffered saline (TBS) plus 0.1% Tween-20 and blotted with primary antibodies (Supplemental Table) overnight at 4°C. After washing with 5% milk in TBS plus 0.1% Tween-20, secondary antibody was applied for one hour using donkey anti-mouse 790 or donkey anti-rabbit 790 (Jackson ImmunoResearch). Blots were washed with TBS and imaged using the Li-COR Odyssey CLx imaging system (LI-COR Biosciences). For the synaptic maturation of RGCs over time, Histone H3 was used as a loading control. Fluorescence intensity was calculated for each band and normalized to Histone H3 at week 1. A fold change was generated by dividing each normalized value for each synaptic protein to the normalized intensity at week 1 (n=3). For RGCs grown alone or with astrocytes, BRN3B was used as a loading control to normalize to RGC-specific protein. Normalized relative fold changes for synaptic proteins were calculated by dividing the BRN3B normalized fluorescence intensity of each of the two groups by the RGCs grown alone (n=3).

### Electrophysiological Recordings

Whole cell patch clamp recordings were made at room temperature (21°C) using HEKA EPC10 amplifier and Patchmaster data acquisition software (HEKA). Recording pipettes with the tip resistance of 4–7 MΩ were fabricated from borosilicate capillary glass (1.2 mm outer diameter, 0.60 mm inner diameter; Sutter Instrument Co.) using a P-1000 puller (Sutter Instrument Co.). The bathing solution contained (in mM): NaCl 140, MgCl_2_ 1, KCl 5, CaCl_2_, HEPES 10, Glucose 10, pH 7.3 (adjusted with NaOH). For current-clamp recordings pipettes were filled with internal solution containing (in mM): KCl 140, MgCl_2_ 5, CaCl_2_ 2.5, EGTA 5, HEPES 10, pH 7.3 (adjusted with KOH) and for voltage-clamp recordings the internal solution contained (in mM): CsF 130, NaCl 10, HEPES 10, CsEGTA (EGTA in CsOH). Inward sodium currents were recorded in voltage-clamp mode with a holding potential of −80mV. Spontaneous AP activity was recorded in current-clamp mode at 0pA immediately after establishing whole-cell configuration. For evoked APs, the cells were allowed to stabilize in the whole-cell configuration for 2 min before initiating either step current injections with incremental steps of 2pA up to 40pA (pulse duration of 500 ms) or ramp current injection with depolarization from 0 to 40 pA for 1s (V_hold_= −80 mV). AP threshold was estimated from the first evoked spike using ramp current injection and was calculated using dV/dt method. AP amplitude was measured as height of the peak from threshold and AP duration was measured as width at the threshold. When necessary, a small bias current was injected to maintain a similar baseline membrane potential (near −70 mV) before depolarizing current injections. Spontaneous postsynaptic currents were measured in the voltage clamp mode at −80mV.

## Author Contributions

K.B.L., experimental design, data collection and interpretation and manuscript writing. R.V., experimental design, data collection and interpretation and manuscript writing. S.K.O., experimental design, data collection. E.F., data collection. M.C.E., experimental design and data interpretation. A.J.B., experimental design and data interpretation. T.C., experimental design and data interpretation. J.S.M., experimental design, data interpretation, manuscript writing, and final approval of manuscript.

## Acknowledgements

We would like to thank Dr. Donald Zack and Dr. Valentin Sluch for providing and sharing the H7 BRN3:tdTomatoThy1.2 hPSC line used in the current study. Grant support was provided by the National Eye Institute (R01 EY024984 to JSM), Indiana Department of Health Brain and Spinal Cord Injury Fund (JSM), an IU Collaborative Research Grant from the Office of the Vice President for Research (JSM), an award from the IU Signature Center for Brain and Spinal Cord Injury (JSM), an IUPUI Graduate Office First Year University Fellowship (KBL), the Purdue Research Foundation Fellowship (KBL), and the Stark Neuroscience Research Institute/Eli Lilly and Company predoctoral fellowship (SKO). This publication was also made possible with partial support from Grant #s UL1TR00108 and UL1TR002529 (A. Shekhar, PI) from the National Institutes of Health, National Center for Advancing Translational Sciences, Clinical and Translational Sciences Award (to SKO and KBL, respectively).

---

**Table S1:** The table represents the information for primary antibodies used throughout the study including the catalog number and their respective dilutions.

| Table S1: Primary Antibody Information |  |  |  |
| --- | --- | --- | --- |
| Primary Antibodies | Company | Catalog Number | Dilution |
| RFP | Rockland | 600-401-379 | 1:2000 |
| GFAP | Millipore | MAB360 | 1:200 |
| GFAP | Cell Signaling Technology | D1F4Q | 1:200 |
| SV2 | DSHB | SV2A | 1:10 |
| Vimentin | Santa Cruz | Sc6260 | 1:100 |
| S100β | AbCam | Ab11178 | 1:200 |
| SOX2 | R&D Systems | AF2018 | 1:1000 |
| EAAT1 | Santa Cruz | Sc515839 | 1:100 |
| Nestin | Santa Cruz | Sc23927 | 1:100 |
| Synapsin-1 | Calbiochem | 574777 | 1:1000 |
| vGlut2 | Cell Signaling Technology | 71555 | 1:1000 |
| SNAP25 | Millipore | MAB331 | 1:500 |
| Neurexin-1 | Millipore | ABN161-I | 1:1000 |
| SynGap | ThermoScientific | Pa1-046 | 1:1000 |
| BRN3B | Santa Cruz | Sc514474 | 1:500 |
| H3 | Cell Signaling Technology | 9715S | 1:1000 |

**Figure S1:**
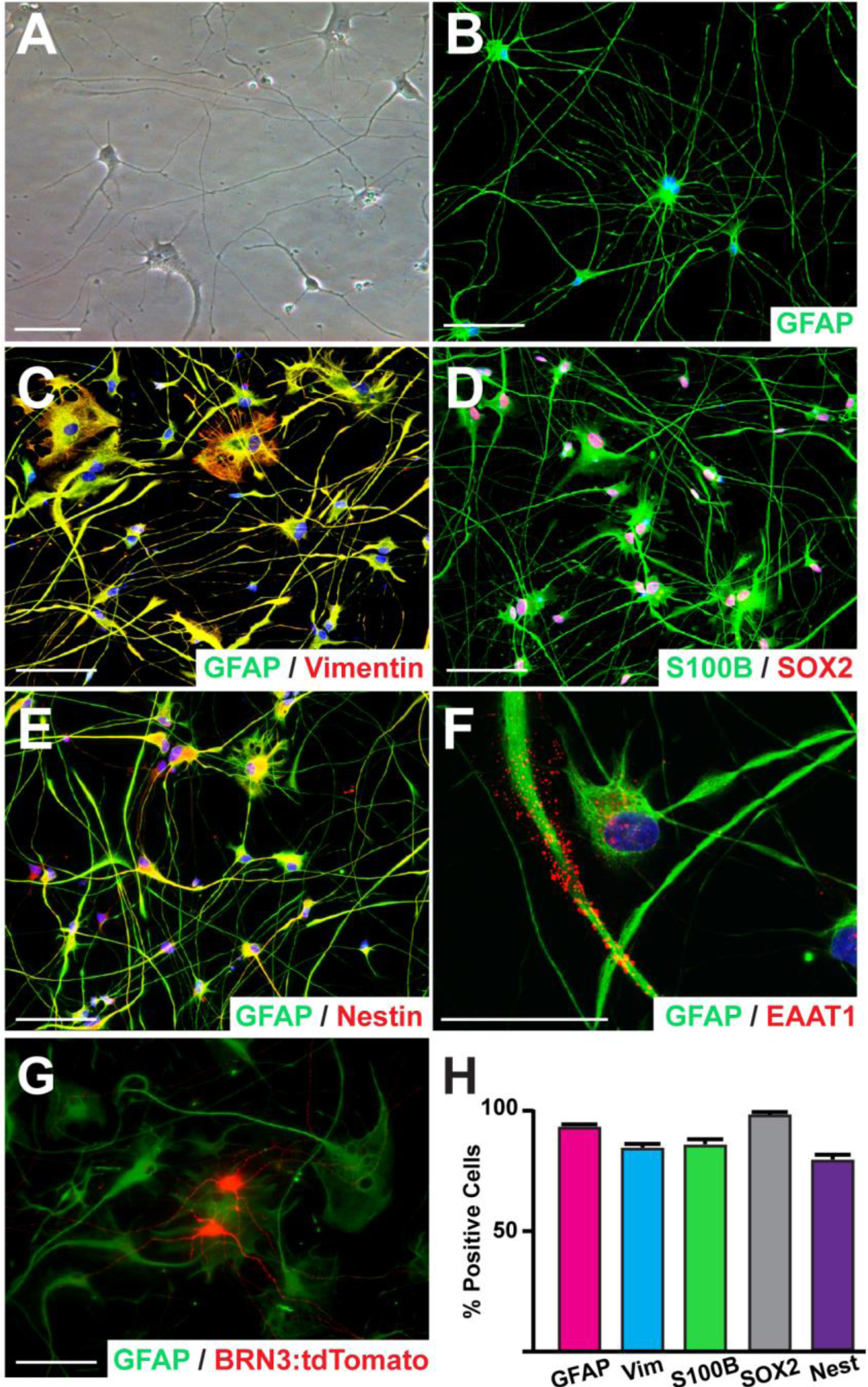
Derivation of astrocytes from hPSCs and establishment of an astrocyte-RGC co-culture system. (A) Bright field microscopy displayed the characteristic morphology of hPSC-derived astrocytes. (B-F) hPSC-derived astrocytes expressed a variety of associated markers including GFAP (B), Vimentin (C), S100β (D), Nestin (E), and EAAT1 (F). (G) tdTomato-expressing hPSC-derived RGCs were co-cultured with GFAP-positive hPSC-derived astrocytes, leading to the extension of long neurites. (H) Quantification of immunocytochemistry demonstrated the percentage of cells expressing the indicated markers within hPSC-derived astrocyte populations. Error bars represent S.E.M (n=3).

**Figure S2:**
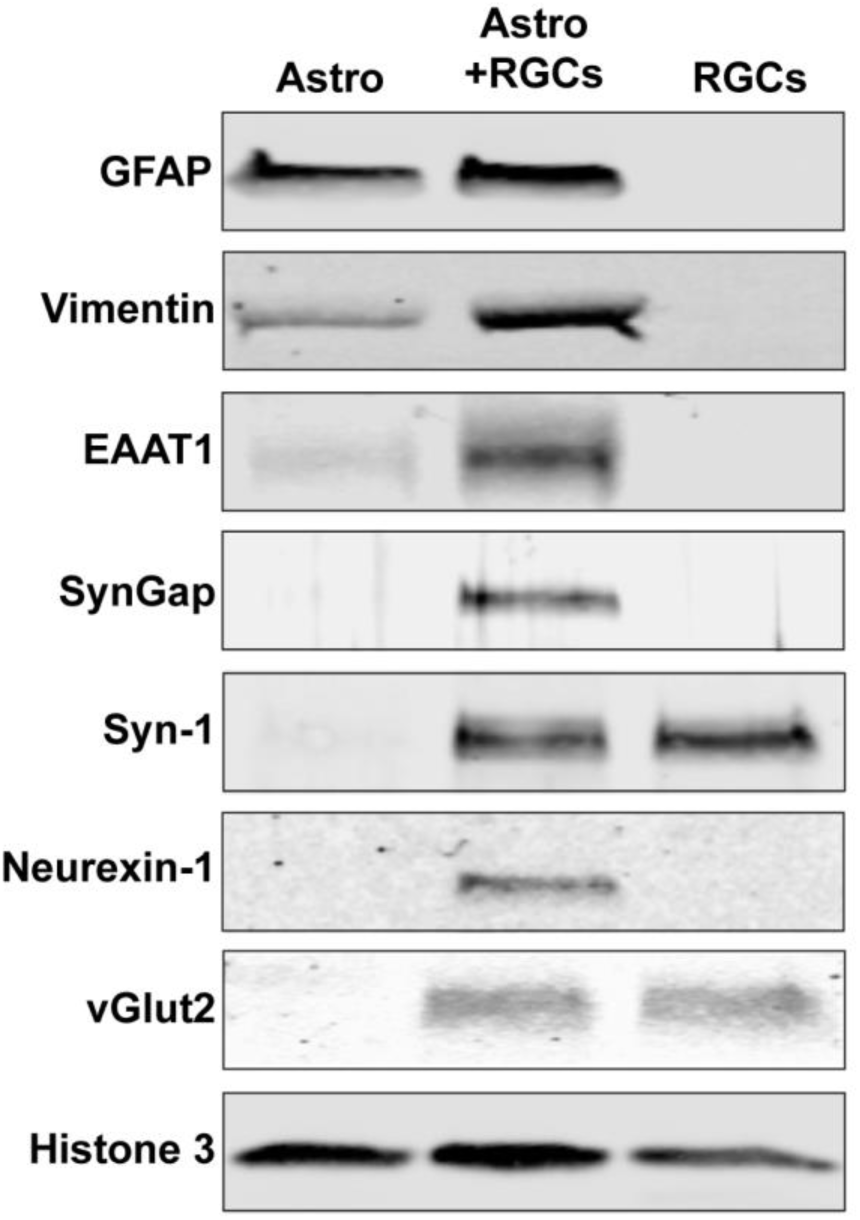
Co-culture of hPSC-derived RGCs with astrocytes leads to the enhanced expression of functional proteins. Western blot analyses demonstrated the expression of astrocyte-associated proteins including GFAP, Vimentin, and EAAT1 in astrocyte cultures and co-cultures, but not in RGCs alone. Similarly, the expression of synaptic proteins including SynGap, Neurexin-1, Synapsin-1, and vGlut2 was observed in RGC cultures and co-cultures, but not astrocytes alone. The co-culture of astrocytes with RGCs led to the expression of synaptic proteins including SynGap and Neurexin-1 that was not observed in RGCs alone, as well as the enhanced expression of EAAT1.

